# The dynamics of starvation and recovery

**DOI:** 10.1101/219519

**Authors:** Justin D. Yeakel, Christopher P. Kempes, Sidney Redner

## Abstract

The eco-evolutionary dynamics of species are fundamentally linked to the energetic constraints of its constituent individuals. Of particular importance is the interplay between reproduction and the dynamics of starvation and recovery. To elucidate this interplay, we introduce a nutritional state-structured model that incorporates two classes of consumer: nutritionally replete, reproducing consumers, and undernourished, non-reproducing consumers. We obtain strong constraints on starvation and recovery rates by deriving allometric scaling relationships and find that population dynamics are typically driven to a steady state. Moreover, these rates fall within a ‘refuge’ in parameter space, where the probability of population extinction is minimized. We also show that our model provides a natural framework to predict maximum mammalian body size by determining the relative stability of an otherwise homogeneous population to a competing population with altered percent body fat. This framework provides a principled mechanism for a selective driver of Cope’s rule.

---

The behavioral ecology of all organisms is influenced by their energetic states, which directly impacts how they invest reserves in uncertain environments. Such behaviors are generally manifested as tradeoffs between investing in somatic maintenance and growth, or allocating energy towards reproduction [1–3]. The timing of these behaviors responds to selective pressure, as the choice of the investment impacts future fitness [4–6]. The influence of resource limitation on an organism’s ability to maintain its nutritional stores may lead to repeated delays or shifts in reproduction over the course of an organism’s life.

The balance between (a) somatic growth and maintenance, and (b) reproduction depends on resource availability [7]. For example, reindeer invest less in calves born after harsh winters (when the mother’s energetic state is depleted) than in calves born after moderate winters [8]. Many bird species invest differently in broods during periods of resource scarcity [9, 10], sometimes delaying or even foregoing reproduction for a breeding season [1, 11, 12]. Even freshwater and marine zooplankton have been observed to avoid reproduction under nutritional stress [13], and those that do reproduce have lower survival rates [2]. Organisms may also separate maintenance and growth from reproduction over space and time: many salmonids, birds, and some mammals return to migratory breeding grounds to reproduce after one or multiple seasons in resource-rich environments where they accumulate reserves [14–16].

Physiology also plays an important role in regulating reproductive expenditures during periods of resource limitation. Many mammals (47 species in 10 families) exhibit delayed implantation, whereby females postpone fetal development until nutritional reserves can be accumulated [17, 18]. Many other species (including humans) suffer irregular menstrual cycling and higher abortion rates during periods of nutritional stress [19, 20]. In the extreme case of unicellular organisms, nutrition directly controls growth to a reproductive state [3, 21]. The existence of so many independently evolved mechanisms across such a diverse suite of organisms highlights the near-universality of the fundamental tradeoff between somatic and reproductive investment.

Including individual energetic dynamics [22] in a population-level framework [22, 23] is challenging [24]. A common simplifying approach is the classic Lotka-Volterra (LV) model, which assumes that consumer population growth rate depends linearly on resource density [25]. Here, we introduce an alternative approach— the Nutritional State-structured Model (NSM)—that accounts for resource limitation via explicit starvation. In contrast to the LV model, the NSM incorporates two consumer states: hungry and full, with only the former susceptible to mortality and only the latter possessing sufficient energetic reserves to reproduce. Additionally, we incorporate allometrically derived constraints on the time scales for reproduction [3], starvation, and recovery. Our model makes several basic predictions: (i) the dynamics are typically driven to a refuge far from cyclic behavior and extinction risk, (ii) the steady-state conditions of the NSM accurately predict the measured biomass densities for mammals described by Damuth’s law [26–29], (iii) there is an allometrically constrained upper-bound for mammalian body size, and (iv) the NSM provides a selective mechanism for the evolution of larger body size, known as Cope’s rule [30–33].

## Nutritional state-structured model (NSM)

We begin by defining the nutritional state-structured population model, where the consumer population is partitioned into two states: (a) an energetically replete (full) state *F*, where the consumer reproduces at a constant rate λ and does not die from starvation, and (b) an energetically deficient (hungry) state *H*, where the consumer does not reproduce but dies by starvation at rate *μ*. The dynamics of the underlying resource *R* are governed by logistic growth with an intrinsic growth rate and a carrying capacity *C*. The rate at which consumers transition between states and consume resources is dependent on their number, the abundance of resources, the efficiency of converting resources into metabolism, and how that metabolism is partitioned between maintenance and growth purposes. We provide a physiologically and energetically mechanistic model for each of these dynamics and constants (see the Supplementary Information (SI)), and show that the system produces a simple non-dimensional form which we describe below.

**Figure 1:**
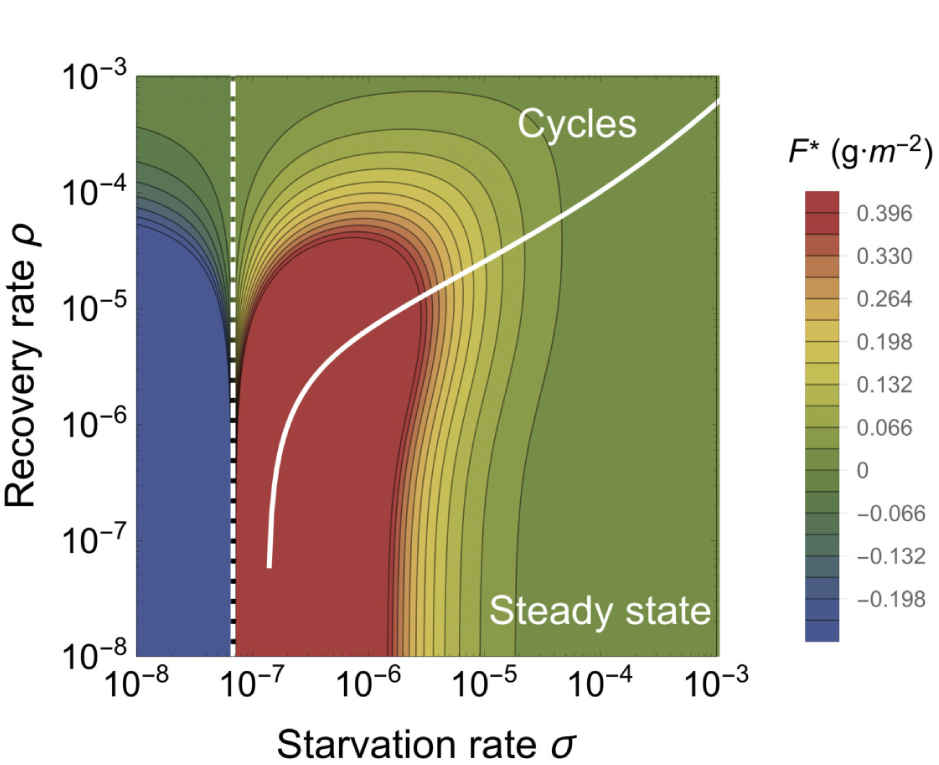
The transcritical (TC; dashed line) and Hopf bifurcation (solid line) as a function of the starvation rate *σ* and recovery rate *ρ* for a 100g consumer. These bifurcation conditions separate parameter space into unphysical (left of the TC), cyclic, and steady state dynamic regimes. The colors show the steady state densities for the energetically replete consumers *F^*^*.

Consumers transition from the full state *F* to the hungry state *H* at a rate σ—the starvation rate—and also in proportion to the absence of resources (1 — *R*) (the maximum resource density has been non dimensionalized to 1; see SI). Conversely, consumers recover from state *H* to state *F* at rate *ξρ* and in proportion to *R*, where *ξ* represents a ratio between maximal resource consumption and the carrying capacity of the resource. The resources that are eaten by hungry consumers (at rate *ρR* + *δ*) account for their somatic growth (*ρR*) and maintenance (*δ*). Full consumers eat resources at a constant rate *β* that accounts for maximal maintenance and somatic growth (see the SI for mechanistic derivations of these rates from resource energetics). The NSM represents an ecologically motivated fundamental extension of the idealized starving random walk model of foraging, which focuses on resource depletion, to include reproduction and resource replenishment [34–36], and is a more general formulation than previous models that incorporate starvation [37].

In the mean-field approximation, in which the consumers and resources are perfectly mixed, their densities are governed by the rate equations

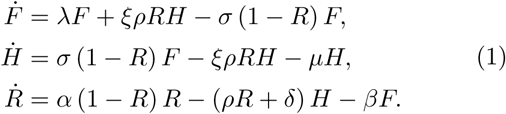

This system of nondimensional equations follows from a set of first-principle relationships for resource consumption and growth (see the SI for a full derivation and the dimensional form). Notice that the total consumer density *F* + *H* evolves according to 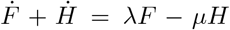. This resembles the equation of motion for the predator density in the **LV** model [38], except that the resource density does not appear in the growth term. The rate of reproduction is independent of resource density because the full consumer partitions a constant amount of energy towards reproduction, whereas a hungry consumer partitions no energy towards reproduction. Similarly, the consumer maintenance terms (*δH* and *βF*) are also independent of resource density because they represent a minimal energetic requirement for consumers in the *H* and *F* state, respectively.

### Steady states of the NSM

From the single internal fixed point (Eq. (2), see Methods), an obvious constraint on the NSM is that the reproduction rate λ must be less than the starvation rate *σ*, so that the consumer and resource densities are positive. The condition *σ* = *λ* represents a transcritical (TC) bifurcation [39] that demarcates a physical from an unphysical (negative steady-state densities) regime. The biological implication of the constraint *λ* < *σ* has a simple interpretation—the rate at which a macroscopic organism loses mass due to lack of resources is generally much faster than the rate of reproduction. As we will discuss below, this inequality is also a natural consequence of allometric constraints [3] for organisms within empirically observed body size ranges.

In the physical regime of *λ* < *σ*, the fixed point (2) may either be a stable node or a limit cycle (Fig. 1). In continuous-time systems, a limit cycle arises when a pair of complex conjugate eigenvalues crosses the imaginary axis to attain positive real parts [40]. This Hopf bifurcation is defined by Det(**S**) = 0, with **S** the Sylvester matrix, which is composed of the coefficients of the characteristic polynomial of the Jacobian matrix [41]. As the system parameters are tuned to be within the stable regime, but close to the Hopf bifurcation, the amplitude of the transient cycles becomes large. Given that ecological systems are constantly being perturbed [42], the onset of transient cycles, even though they decay with time in the mean-field description, can increase extinction risk [43–45].

When the starvation rate *σ* ≫ *λ*, a substantial fraction of the consumers are driven to the hungry non-reproducing state. Because reproduction is inhibited, there is a low steady-state consumer density and a high steady-state resource density. However, if *σ*/*λ* → 1 from above, the population is overloaded with energetically-replete (reproducing) individuals, thereby promoting transient oscillations between the consumer and resource densities (Fig. 1). If the starvation rate is low enough that the Hopf bifurcation is crossed, these oscillations become stable. This threshold occurs at higher values of the starvation rate as the recovery rate *ρ* increases, such that the range of parameter space giving rise to cyclic dynamics also increases with higher recovery rates.

**Figure 2:**
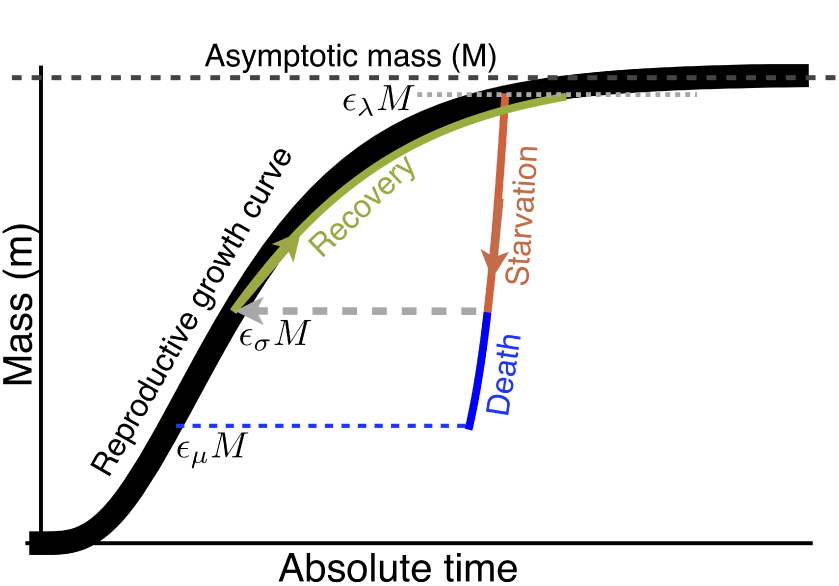
The growth trajectory over absolute time of an individual organism as a function of body mass. Initial growth follows the black trajectory to an energetically replete reproductive adult mass of *m* = *∊_λ_M* (see Methods). Starvation follows the red trajectory to *m* = *∊_σ_∊_λ_M*. Recovery follows the green curve to the replete adult mass, where this trajectory differs from the original growth because only fat is being regrown which requires a longer time to reach *∊_λ_M*. Alternatively, death from starvation follows the blue trajectory to *m* = *∊_μ_∊_λ_M*.

## Results

### The allometry of extinction risk

While there are no *a priori* constraints on the parameters in the NSM, we expect that each species should be restricted to a distinct portion of the parameter space. We use allometric scaling relations to constrain the covariation of rates in a principled and biologically meaningful manner (see Methods). Allometric scaling relations highlight common constraints and average trends across large ranges in body size and species diversity. Many of these relations can be derived from a small set of assumptions. In the Methods we describe our framework to determine the covariation of timescales and rates across a range of body sizes for each of the key parameters of our model (cf. Ref. [46]).

Nearly all of the rates described in the NSM are determined by consumer metabolism, which can be used to describe a variety of organismal features [47]. We derive, from first principles, the relationships for the rates of reproduction, starvation, recovery, and mortality as a function of an organism’s body size and metabolic rate (see Methods). Because we aim to explore the starvation-recovery dynamics as a function of an organism’s body mass *M*, we parameterize these rates in terms of the *percent* gain and loss of the asymptotic (maximum) body mass, *∊M*, where different values of *∊* define different states of the consumer (Fig. 2; see Methods for derivations of allometrically constrained rate equations). Although the rate equations (1) are general and can in principle be used to explore the starvation recovery dynamics for most organisms, here we focus on allometric relationships for terrestrial-bound lower-trophic level endotherms (see the SI for values), specifically herbivorous mammals, which range from a minimum of *M* ≈ 1g (the Etruscan shrew *Suncus etruscus*) to a maximum of *M* ≈ 10^7^g (the early Oligocene Indricotheriinae and the Miocene Deinotheriinae). Investigating other classes of organisms would simply involve altering the metabolic exponents and scalings associated with *∊*. Moreover, we emphasize that our allometric equations (see Methods) describe mean relationships, and do not account for the (sometimes considerable) variance associated with individual species. We note that including additional allometrically-scaled mortality terms to both *F* and *H* does not change the form of our model nor impact our quantitative findings (see SI for the derivation).

**Figure 3:**
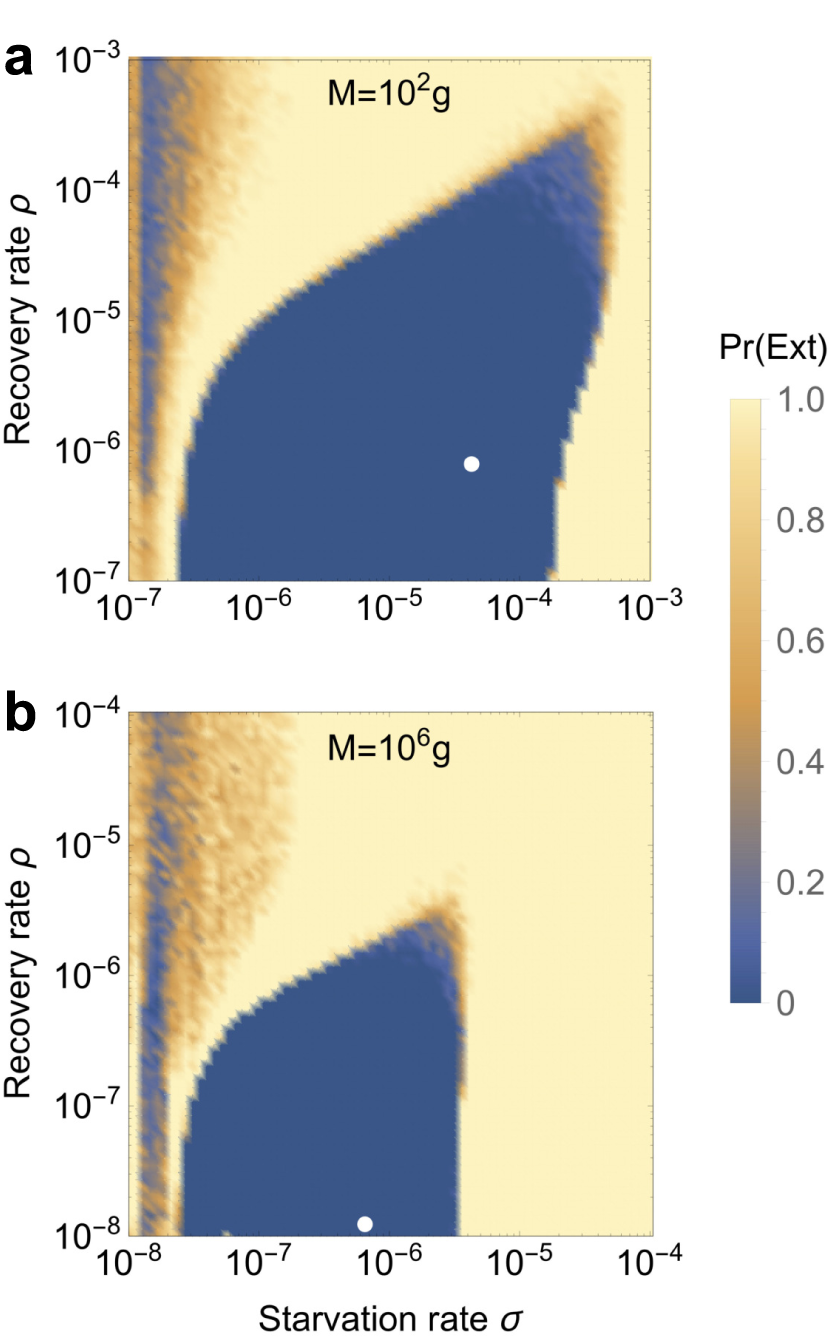
Probability of extinction for a consumer with (**a**) *M* = 10^2^g and (**b**) *M* = 10^6^g as a function of the starvation rate *σ* and recovery rate *ρ*, where the initial density is given as (*XF^*^*, *XH^*^*, *R^*^*), where *X* is a random uniform variable in [0, 2]. Note the change in scale in panel **b.** Extinction is defined as the population trajectory falling below 0.2× the allometrically constrained steady state. The white points denote the allometrically constrained starvation and recovery rate.

As the allometric derivations of the NSM rate laws reveal (see Methods), starvation and recovery rates are not independent parameters, and the biologically relevant portion of the phase space shown in Fig. 1 is constrained via covarying parameters. Given the parameters of terrestrial endotherms, we find that the starvation rate *σ* and the recovery rate *ρ* are constrained to lie within a small region of potential values for the known range of body sizes *M*. Indeed, starvation and recovery rates across all values of *M* fall squarely in the steady-state region at some distance from the Hopf bifurcation. This suggests that cyclic population dynamics should be rare, particularly in resource-limited environments.

Higher rates of starvation result in a larger flux of the population to the hungry state. In this state, reproduction is absent, thus increasing the likelihood of extinction. From the perspective of population survival, it is the rate of starvation relative to the rate of recovery that determines the long-term dynamics of the various species (Fig. 1). We therefore examine the competing effects of cyclic dynamics vs. changes in steady-state density on extinction risk, both as functions of *σ* and *ρ*. To this end, we computed the probability of extinction, where we define extinction as a population trajectory falling below one fifth of the allometrically constrained steady state at any time between *t* = 10^8^ and *t* = 10^10^. This procedure was repeated for 50 replicates of the continuous-time system shown in Eq. 1 for organisms with mass ranging from 10^2^ to 10^6^ grams. In each replicate the initial densities were chosen to be (*XF^*^*, *XH^*^*, *R^*^*), with *X* a random variable uniformly distributed in [0, 2]. By allowing the rate of starvation to vary, we assessed extinction risk across a range of values for *σ* and *ρ* between ca. 10^−8^ to 10^−3^. Higher rates of extinction correspond to both large *σ* if *ρ* is small, and large *ρ* if *σ* is small. In the former case, increased extinction risk arises because of the decrease in the steady-state consumer population density (Figs. 1b, 3). In the latter case, the increased extinction risk results from higher-amplitude transient cycles as the system nears the Hopf bifurcation (Fig. 3). This interplay creates an ‘extinction refuge’, such that for a constrained range of *σ* and *ρ,* extinction probabilities are minimized.

We find that the allometrically constrained values of *σ* and *ρ*, each representing different trajectories along the ontogenetic curve (Fig. 2), fall squarely within the extinction refuge across a range of *M* (Fig. 3a,b, white points). These values are close enough to the Hopf bifurcation to avoid low steady-state densities, yet distant enough to avoid large-amplitude transient cycles. Allometric values of *σ* and *ρ* fall within this relatively small window, which supports the possibility that a selective mechanism has constrained the physiological conditions driving starvation and recovery rates within populations. Such a mechanism would select for organism physiology that generates appropriate *σ* and *ρ* values that minimize extinction risk. This selection could occur via the tuning of body fat percentages, metabolic rates, and/or biomass maintenance efficiencies. We also find that as body size increases, the size of the low extinction-risk parameter space shrinks (Fig. 3b), suggesting that the population dynamics for larger organisms are more sensitive to variability in physiological rates. This finding is in accordance with, and may serve as contributing support for, observations of increased extinction risk among larger mammals [48].

**Figure 4:**
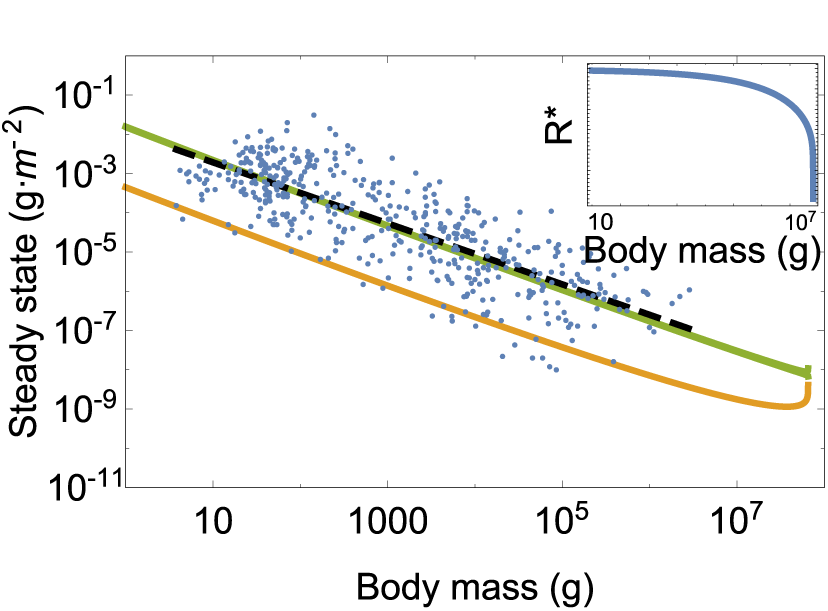
Consumer steady states *F^*^* (green) and *H^*^* (orange) as a function of body mass along with the data from Damuth [26]. Inset: Resource steady state *R^*^* as a function of consumer body mass.

### Damuth’s Law and body size limits

The NSM correctly predicts that smaller species have larger steady-state population densities (Fig. 4). Similar predictions have been made for carnivore populations using alternative consumer-resource models [49]. Moreover, we show that the NSM provides independent theoretical support for Damuth’s Law [26–29]. Damuth’s law shows that species abundances, *N^*^*, follow *N^*^* = 0.01M^−0.78^ (g m^−2^). Figure 4 shows that both *F*^*^ and *H^*^* scale as *M^−η^*, with *η* ≈ 3/4, over a wide range of organismal sizes and that *F^*^* + *H^*^* closely matches the best fit to Damuth’s data. Remarkably, this result illustrates that the steady state values of the NSM combined with the derived timescales naturally give rise to Damuth’s law. While the previous metabolic studies supporting Damuth’s law provided arguments for the value of the exponent [27], these studies are only able to infer the normalization constant (0.01 g^1.78^ m^−2^ in the above equation) from the data (see SI for a discussion of the energy equivalence hypothesis related to these metabolic arguments). Our model predicts not only the exponent but also the normalization constant by explicitly including the resource dynamics and the parameters that determine growth and consumption. It should be noted that density relationships of individual clades follow a more shallow scaling relationship than predicted by Damuth’s law [29]. In the context of our model, this finding suggests that future work may be able to anticipate these shifts by accounting for differences in the physiological parameters associated with each clade.

With respect to predicted steady state densities, the total metabolic rate of *F* and *H* becomes infinite at a finite mass, and occurs at the same scale where the steady state resources vanish (Fig. 4). This asymptotic behavior is governed by body sizes at which *∊_μ_* and *∊_λ_* (see Fig. 2) equal zero, causing the timescales (Eqn. 4) to become infinite and the rates *ρ* and *σ* to equal zero. The *μ* = 0 asymptote occurs first when *f*_0_*M*^*γ*−1^ + *u*_0_*M*^*ζ*−1^ = 1, and corresponds to (*F^*^*, *H^*^*, *R^*^*) = (0, 0, 0). This point predicts an upper bound on mammalian body size at *M*_max_ = 6.54 × 10^7^ (g). Moreover, *M*_max_, which is entirely determined by the population-level consequences of energetic constraints, is within an order of magnitude of the maximum body size observed in the North American mammalian fossil record [30], as well as the mass predicted from an evolutionary model of body size evolution [31]. We emphasize that the asymptotic behavior and predicted upper bound depend only on the scaling of body composition and are independent of the resource parameters. The prediction of an asymptotic limit on mammalian size parallels work on microbial life where an upper and lower bound on bacterial size, and an upper bound on single cell eukaryotic size, is predicted from similar growth and energetic scaling relationships [3, 50]. It has also been shown that models that incorporate the allometry of hunting and resting combined with foraging time predicts a maximum carnivore size between 7 × 10^5^ and 1.1 × 10^6^ (g) [51, 52]. Similarly, the maximum body size within a particular lineage has been shown to scale with the metabolic normalization constant [53]. This complementary approach is based on the balance between growth and mortality, and suggests that future connections between the scaling of fat and muscle mass should systematically be connected with *B*_0_ when comparing lineages.

### A mechanism for Cope’s rule

Metabolite transport constraints are widely thought to place strict boundaries on biological scaling [47, 54, 55] and thereby lead to specific predictions on the minimum possible body size for organisms [56]. Above this bound, a number of energetic and evolutionary mechanisms have been explored to assess the costs and benefits associated with larger body masses, particularly for mammals. One important such example is the *fasting endurance hypothesis*, which contends that larger body size, with consequent lower metabolic rates and increased ability to maintain more endogenous energetic reserves, may buffer organisms against environmental fluctuations in resource availability [57]. Over evolutionary time, terrestrial mammalian lineages show a significant trend towards larger body size—Cope’s rule [30–33]. It is thought that within-lineage drivers generate selection towards an optimal upper bound of roughly 10^7^ (g) [30], a value that is likely limited by higher extinction risk for large taxa over longer timescales [31]. These trends are thought to be driven by a combination of climate change and niche availability [33]; however the underpinning energetic costs and benefits of larger body sizes, and how they influence dynamics over ecological timescales, have not been explored.

The NSM predicts that the steady state resource density *R^*^* decreases with increasing body size of the consumer population (Fig. 4, inset), and classic resource competition theory predicts that the species surviving on the lowest resource abundance will outcompete others [58–60]. Thus, the combined NSM steady-state dynamics and allometric timescales (see Eq. (4)) predict that larger mammals have an intrinsic competitive advantage given a common resource.

**Figure 5:**
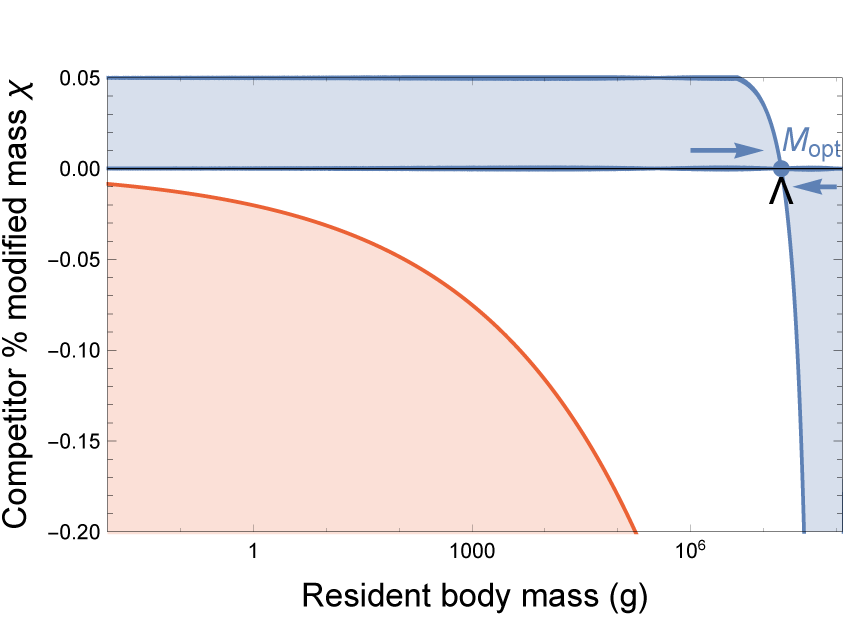
Competitive outcomes for a resident species with body mass *M* vs. a closely related competing species with modified body mass *M′* = *M*(1 + *χ*). The blue region denotes proportions of modified mass *χ* resulting in exclusion of the resident species. The red region denotes values of *χ* that result in a mass that is below the starvation threshold and are thus infeasible. Arrows point to the predicted optimal mass from our model *M*_opt_ = 1:748 × 10^7^, which may serve as an evolutionary attractor for body mass. The black wedge points to the largest body mass known for terrestrial mammals (*Deinotherium* spp.) at 1:74 × 10^7^ (g) [32].

However, the above resource relationships do not offer a mechanism for how body size is selected. We directly assess competitive outcome between two closely related species: a resident species of mass *M*, and a competing species (denoted by ′) where individuals have a different proportion of body fat such that *M*′ = *M*(1 + *χ*). For *χ* < 0, the competing individuals have fewer metabolic reserves than the resident species and vice versa for *χ* > 0. For the allowable values of *χ* (see SI), the mass of the competitor *M*′ should exceed the minimal amount of body fat, 1 + *χ* > *∊_σ_*, and the adjusted time to reproduce must be positive, which, given Eq. 4, implies that 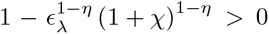. These conditions imply that 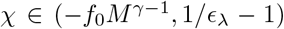 where the upper bound approximately equals 0. 05 and the lower bound is mass-dependent. The modified mass of the competitor leads to altered rates of starvation *σ*(*M*′), recovery *ρ*(*M*′), and the maintenance of both starving *δ*(*M*′) and full consumers *β*(*M*′) (see the SI for derivations of competitor rates). Importantly, *∊_σ_*, which determines the point along the growth curve that defines the body composition of starved foragers, is assumed to remain unchanged for the competing population (see SI).

To assess the susceptibility of the resident species to competitive exclusion, we determine which consumer pushes the steady-state resource density *R^*^* to lower values for a given value of *χ*, with the expectation that a population capable of surviving on lower resource densities has a competitive advantage [58]. We find that for *M* ≤ 1.748 × 10^7^ (g), having additional body fat (*χ* > 0) results in a lower steady state resource density (*R*′^*^ < *R*^*^), such that the competitor has an intrinsic advantage over the resident species (Fig. 5). However, for *M* > 1.748 × 10^7^ (g), leaner individuals (*χ* < 0) have lower resource steady state densities.

The observed switch in susceptibility as a function of *χ* at *M*_opt_ = 1.748 × 10^7^ (g) thus serves as an attractor, such that the NSM predicts organismal mass to increase if *M* < *M*_opt_ and decrease if *M* > *M*_opt_. This value is close to but smaller than the asymptotic upper bound for terrestrial mammal body size predicted by the NSM, and is remarkably close to independent estimates of the largest land mammals, the early Oligocene *Indricotherium* at ≈ 1.5 × 10^7^ (g) and the late Miocene *Deinotherium* at ≈ 1. 74 × 10^7^ (g) [32]. Additionally, our calculation of *M*_opt_ as a function of mass-dependent physiological rates is similar to theoretical estimates of maximum body size [31], and provides independent theoretical support for the observation of a ‘maximum body size attractor’ explored by Alroy [30].

An optimal size for mammals at intermediate body mass was predicted by Brown et al. based on reproductive maximization and the transition between hungry and full individuals [54]. By coupling the NSM to resource dynamics as well as introducing an explicit treatment of storage, we show that species with larger body masses have an inherent competitive advantage for size classes up to *M*_opt_ = 1.748 × 10^7^ based on resource competition. Moreover, the mass distributions in Ref. [54] show that intermediate mammal sizes have the greatest species diversity, in contrast to our efforts, which consider total biomass and predict a much larger *M_opt_*. Compellingly, recent work shows that many communities can be dominated by the biomass of the large [61]. While the state of the environment as well as the competitive landscape will determine whether specific body sizes are selected for or against [33], we propose that the dynamics of starvation and recovery described in the NSM provide a general selective mechanism for the evolution of larger body size among terrestrial mammals.

## Discussion

The energetics associated with somatic maintenance, growth, and reproduction are important elements that influence the dynamics of all populations [11]. The NSM incorporates the dynamics of starvation and recovery that are expected to occur in resource-limited environments. We found that incorporating allometrically-determined rates into the NSM predicts that: (i) extinction risk is minimized, (ii) the derived steady-states quantitatively reproduce Damuth’s law, and (iii) the selective mechanism for the evolution of larger body sizes agrees with Cope’s rule. The NSM offers a means by which the dynamic consequences of energetic constraints can be assessed using macroscale interactions between and among species.

## Methods

### Analytical solution to the NSM

Equation (1) has three fixed points: two trivial fixed points at (*F^*^*, *H^*^*, *R^*^*) = (0, 0, 0) and (0, 0,1), and one non-trivial, internal fixed point at

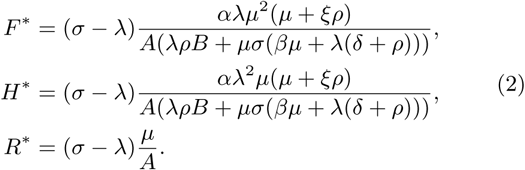

where *A* = (*λξρ* + *μσ*) and *B* = (*βμξ* + *δλξ* − *λμ*). The stability of this fixed point is determined by the Jacobian matrix **J**, with *J_ij_* = *∂X_i_*/*∂X_j_*, when evaluated at the internal fixed point, and **X** is the vector (*F, H, R*). The parameters in Eq. (1) are such that the real part of the largest eigenvalue of **J** is negative, so that the system is stable with respect to small perturbations from the fixed point. Because this fixed point is unique, it is the global attractor for all population trajectories for any initial condition where the resource and consumer densities are both nonzero.

### Metabolic scaling relationships

The scaling relation between an organism’s metabolic rate *B* and its body mass *M* at reproductive maturity is known to scale as *B* = *B*_0_*M^η^*, where the scaling exponent *η* is typically close to 2/3 or 3/4 for metazoans (e.g., Ref. [47, 62]), and has taxonomic shifts for unicellular species between *η* ≈ 1 in eukaryotes and *η* ≈ 1.76 in bacteria [3, 63].

Several efforts have shown how a partitioning of *B* between growth and maintenance purposes can be used to derive a general equation for both the growth trajectories and growth rates of organisms ranging from bacteria to metazoans [3, 64–68]. This relation is derived from the simple balance condition 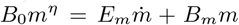, [3, 64–68] where *E_m_* is the energy needed to synthesize a unit of mass, *B_m_* is the metabolic rate to support an existing unit of mass, and *m* is the mass of the organism at any point in its development. This balance has the general solution [3, 69]

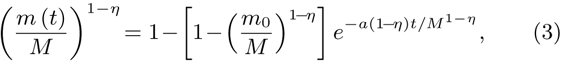

where, for *η* < 1, *M* = (*B*_0_/*B_m_*)^1/(1−*η*)^ is the asymptotic mass, *α* = B_0_/E_m_, and m_0_ is mass at birth, itself varying allometrically (see the SI). We now use this solution to define the timescale for reproduction and recovery from starvation (Fig. 2; see [65] for a detailed presentation of these timescales). The time that an organism takes to reach a particular mass *∊M* is given by the timescale

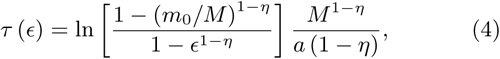

where we define values of e below to describe a variety of timescales, along with the rates related to *τ*. For example, the rate of reproduction is given by the timescale to go from the birth mass to the adult mass. The time to reproduce is given by Equation 4 as *t_λ_* = *τ* (*∊_λ_*), where *∊_λ_* is the fraction of the asymptotic mass where an organism is reproductively mature and should be close to one (typically *∊_λ_* ≈ 0.95 [64]). Our reproductive rate, *λ*, is a specific rate, or the number of offspring produced per time per individual, defined as 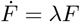. In isolation this functional form gives the population growth *F*(*t*) = *F*_0_*e^λt^* which can be related to the reproductive timescale by assuming that when *t* = *t_λ_* it is also the case that *F* = *νF*_0_, where *ν* − 1 is the number of offspring produced per reproductive cycle. Following this relationship the growth rate is given by *λ* = ln (*ν*) /*t_λ_*, which is the standard relationship (e.g., [68]) and will scales as λ ∝ *M*^*η*−1^ for *M* ≫ *m*_0_ for any constant value of *∊_λ_* [3, 64–67].

The rate of recovery *ρ* = 1/*t_ρ_* requires that an organism accrues sufficient tissue to transition from the hungry to the full state. Since only certain tissues can be digested for energy (for example the brain cannot be degraded to fuel metabolism), we define the rates for starvation, death, and recovery by the timescales required to reach, or return from, specific fractions of the replete-state mass (see the SI, Table I, for parameterizations). We define *m_σ_* = *∊_σ_M*, where *∊_σ_* < 1 is the fraction of replete-state mass where reproduction ceases. This fraction will deviate from a constant if tissue composition systematically scales with adult mass. For example, making use of the observation that body fat in mammals scales with overall body size according to *M*_fat_ = *f*_0_*M^γ^* and assuming that once this mass is fully digested the organism starves, this would imply that *∊_σ_* = 1 − *f*_0_*M^γ^*/*M*. It follows that the recovery timescale, *t_ρ_*, is the time to go from mass *m* = *∊_σ_∊_λ_M* to *m* = *∊_λ_M* (Fig. 2). Using Eqs. (3) and (4) this timescale is given by simply considering the growth curve starting from a mass of 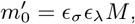, in which case

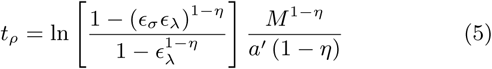

where 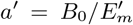 accounts for possible deviations in the biosynthetic energetics during recovery (see the SI). It should be noted that more complicated ontogenetic models explicitly handle storage [67], whereas this feature is implicitly covered by the body fat scaling in our framework.

To determine the starvation rate, σ, we are interested in the time required for an organism to go from a mature adult that reproduces at rate λ, to a reduced-mass hungry state where reproduction is impossible. For starving individuals we assume that an organism must meet its maintenance requirements by using the digestion of existing mass as the sole energy source. This assumption implies the metabolic balance 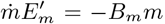 or 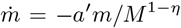 where 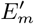 is the amount of energy stored in a unit of existing body mass, which differs from *E_m_*, the energy required to synthesis a unit of biomass [67]. Given the replete mass, *M*, of an organism, the above energy balance prescribes the mass trajectory of a non-consuming organism: 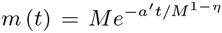. The timescale for starvation is given by the time it takes *m*(*t*) to reach *∊_σ_M*, which gives

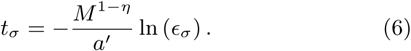

The starvation rate is then σ = 1/*t_σ_*, which scales with replete-state mass as 1/*M*^1−*η*^ ln (1 − *f*_0_*M^γ^*−/*M*). An important feature is that σ does not have a simple scaling dependence on A, which is important for the dynamics that we later discuss.

The time to death should follow a similar relation, but defined by a lower fraction of replete-state mass, *m_μ_* = *∊_μ_M* where *∊_μ_* < *∊_σ_*. Suppose, for example, that an organism dies once it has digested all fat and muscle tissues, and that muscle tissue scales with body mass according to *M*_musc_ = *u*_0_*M^ζ^*. This gives *∊_μ_* = 1 − (*f*_0_*M^γ^* + *u*_0_*M^ζ^*) /*M*. Muscle mass has been shown to be roughly proportional to body mass [70] in mammals and thus *∊_μ_* is merely *∊_μ_* minus a constant. The time to go from starvation to death is the total time to reach *∊_μ_M* minus the time to starve, or *t_μ_* = −*M*^1−*η*^ ln (*∊_μ_*) /*a′* − *t_σ_*, and *μ* = 1/*t_μ_*.

## Supporting Information for “The dynamics of starvation and recovery”

### Mechanisms of Starvation and Recovery

To understand the dynamics of starvation, recovery, reproduction, and resource competition, our framework partitions consumers into two classes: (a) a full class that is able to reproduce and, (b) a hungry class that experiences mortality at a given rate and is unable to reproduce. For the dynamics of growth, reproduction, and resource consumption, past efforts have combined the overall metabolic rate, as dictated by body size, with a growth rate that is dependent on resource abundance and, in turn, dictates resource consumption (see Refs. [1, 2] for a brief review of this perspective). This approach has been used to understand a range of phenomena including a derivation of ontogenetic growth curves from a partitioning of metabolism into maintenance and biosynthesis (e.g. [1, 3–5]) and predictions for the steady-state resource abundance in communities of cells [2]. Here we leverage these mechanisms, combined with several additional concepts, to define our Nutritional State Model (NSM).

We consider the following generalized set of explicit dynamics for starvation, recovery, reproduction, and resource growth and consumption

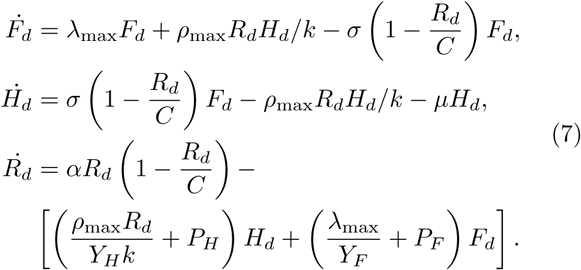

where each term has a mechanistic meaning that we detail below (we will denote the dimensional equations with the subscript _*d*_ before introducing the non-dimensional form that is presented in the main text). In the above equations *Y* represents the yield coefficient (e.g., Refs. [6, 7]) which is the quantity of resources required to build a unit of organism (gram of mammal produced per gram of resource consumed) and *P* is the specific maintenance rate of resource consumption (g resource ⋅ s^−1^ ⋅ g organism^−1^). If we pick *F_d_* and *H_d_* to have units of (g organisms ⋅ m^−2^), then all of the terms of 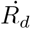, such as 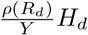, have units of (g resource ⋅ m^−2^ ⋅ s^−1^) which are the units of net primary productivity (**NPP**), a natural choice for 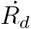. This choice also gives *R_d_* as (g ⋅ m^−2^) which is also a natural unit and is simply the biomass density. In these units *α* (s^−1^) is the specific growth rate of *R_d_*, *C* is the carrying capacity, or maximum density, of *R_d_* in a particular environment, and *k* is the half-saturation constant (half the density of resources that would lead to maximum growth).

We can formally non-dimensionalize this system by the rescaling of *F* = *fF_d_*, *H* = *fH_d_*, *R* = *_q_R_d_*, *t* = *st_d_*, in which case our system of equations becomes

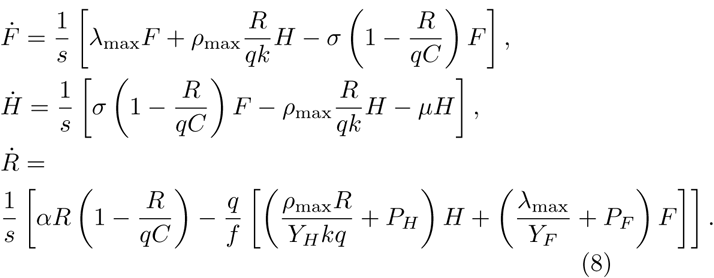

If we make the natural choice of *s* = 1, *q* = 1/*C*, and *f* = 1/*Y_H_ k*, then we are left with

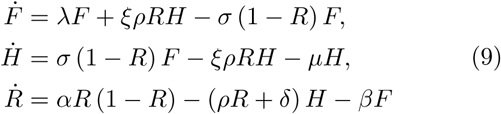

where we have dropped the subscripts on *λ*_max_ and *ρ*_max_ for simplicity, and *ξ* ≡ *C/k*, *δ* ≡ *Y_H_kP_H_*/*C*, and 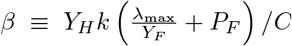. The above equations represent the system of equations presented in the main text.

### Parameter Values and Estimates

All of the parameter values employed in our model have either been directly measured in previous studies or can be estimated from combining several previous studies. Below we outline previous measurements and simple estimates of the parameters.

Metabolic rate has been generally reported to follow an exponent close to *η* = 0.75 (e.g., Refs. [3, 4] and the supplement for Ref. [5]). We make this assumption in the current paper, although alternate exponents, which are known to vary between roughly 0.25 and 1.5 for single species [4], could be easily incorporated into our framework, and this variation is effectively handled by the 20% variations that we consider around mean trends. The exponent not only defines several scalings in our framework, but also the value of the metabolic normalization constant, *B*_0_, given a set of data. For mammals the metabolic normalization constant has been reported to vary between 0.018 (W g^−0.75^) and 0.047 (W g^−0.75^; Refs. [3, 5], where the former value represents basal metabolic rate and the latter represents the field metabolic rate. We employ the field metabolic rate for our NSM model which is appropriate for active mammals (Table 1).

An important feature of our framework is the starting size, *m*_0_, of a mammal which adjusts the overall timescales for reproduction. This starting size is known to follow an allometric relationship with adult mass of the form *m*_0_ = *n*_0_*M^v^* where estimates for the exponent range between 0.71 and 0.94 (see Ref. [8] for a review). We use *m*_0_ = 0.097M^0-92^ [9] which encompasses the widest range of body sizes [8].

The energy to synthesize a unit of biomass, *E_m_*, has been reported to vary between 1800 to 9500 (J g^−1^) (e.g. Refs. [3–5]) in mammals with a mean value across many taxonomic groups of 5, 774 (J g^−1^) [4]. The unit energy available during starvation, *E′*, could range between 7000 (J g^−1^), the return of the total energy stored during ontogeny [5] to a biochemical upper bound of *E’* = 36,000 (J g^−1^) for the energetics of palmitate [5, 10]. For our calculations we use the measured value for bulk tissues of 7000 which assumes that the energy stored during ontogeny is returned during starvation [5].

For the scaling of body composition it has been shown that fat mass follows *M*_fat_ = *f*_0_*M*^*γ*^, with measured relationships following 0.0 8*M*^1.25^ [11], 0.02*M*^1.19^ [12], and 0.026*M*^1.14^ [13]. We use the values from [12] which falls in the middle of this range. Similarly, the muscle mass follows *M*_musc_ = *u*_0_*M^ζ^* with *u*_0_ = 0.383 and *ζ* = 1.00 [13].

Typically the value of *ξ* = *C*/*k* should roughly be 2. The value of *ρ*, *λ*, *σ*, and *μ* are all simple rates (note that we have not rescaled time in our non-dimensionalization) as defined in the maintext. Given that our model considers transitions over entire stages of ontogeny or nutritional states, the value of *Y* must represent yields integrated over entire life stages. Given an energy density of *E_d_* = 18200 (J g^−1^) for grass [14] the maintenance value is given by *P_F_* = *B*_0_*M*^3/4^/*ME_d_*, and the yield for a full organism will be given by *Y_F_* = *ME_d_*/*B_λ_* (g individual ⋅ g grass ^−1^), where *B_λ_* is the lifetime energy use for reaching maturity given by

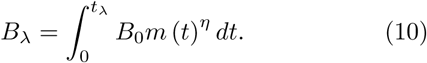

Similarly, the maintenance resource consumption rate for hungry individuals is *P_H_* = *B*_0_(∊_σ_*M*)^3/4^/(∊_σ_*M*)*E_d_*, and the yield for hungry individuals (representing the cost on resources to return to the full state) is given by *Y_H_* = *ME_d_*/*B_ρ_* where

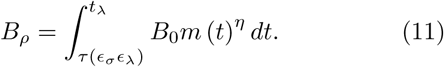

Taken together, these relationships allow us to calculate *ρ, δ*, and *β*.

Finally, the value of *α* can be roughly estimated by the NPP divided by the corresponding biomass densities. From the data in Ref. [15] we estimate the value of *α* to range between 2.81 × 10^−10^ (s^−1^) and 2.19 × 10^−8^ (s^−1^) globally. It should be noted that the value of *α* sets the overall scale of the *F^*^* and *H^*^* steady states along with *B_tot_* for each type. As such, we use *α* as our fit parameter to match these steady states with the data from Damuth [16]. We find that the best fit is *α* = 9.45 × 10^−9^ (s^−1^) which compares well with the calculated range above. However, two points are important to note here: first, our framework predicts the overall scaling of *F^*^* and *H^*^* independently of *α* and this correctly matches data, and second, both the asymptotic behavior and slope of *F^*^* and *H^*^* are independent of *α,* such that our prediction of the maximum mammal size does not depend on *α*.

**Table S1:**
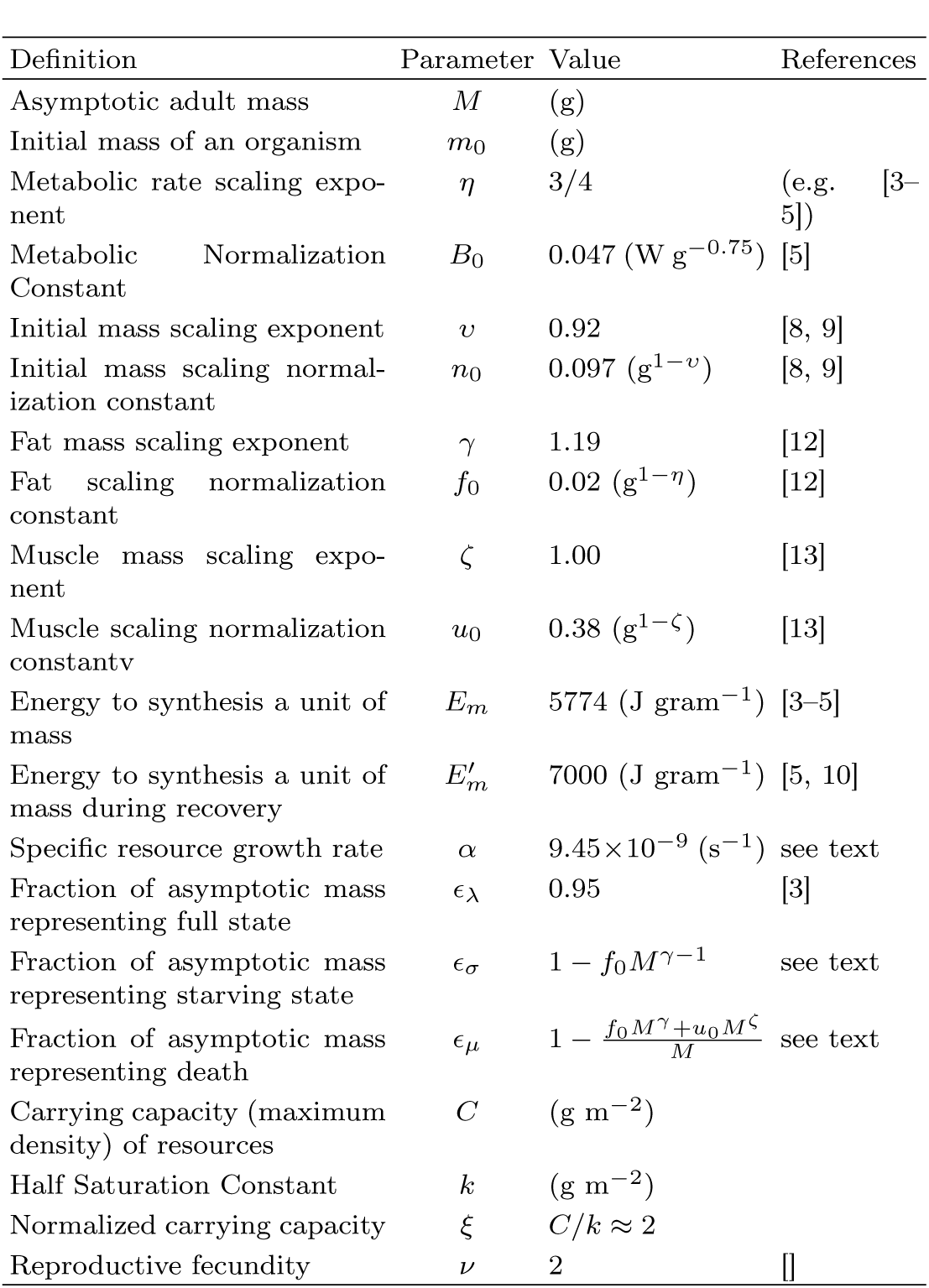
Parameter values for mammals

### Rate equations for invaders with modified body mass

We allow an invading subset of the resident population with mass *M* to have an altered mass *M*′ = *M*(1+*χ*) where *χ* varies between *χ_min_* < 0 and *χ_max_* > 0, where *χ* < 0 denotes a leaner invader and *χ* > 0 denotes an invader with additional reserves of body fat. Importantly, we assume that the invading and resident individuals have the same proportion of non-fat tissues. For the allowable values of *χ* the adjusted mass should exceed the amount of body fat, 1 + *χ* > ∊_σ_, and the adjusted time to reproduce must be positive, which given our solution for *τ*(*∊*) (see main text), implies that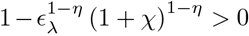. Together these conditions imply that *χ* ∈ (−*f*_0_*M*^γ−1^,1/∊_λ_ − 1) where the upper bound approximately equals 0.05.

Although the starved state of invading organisms remains unchanged, the rate of starvation from the modified full state to the starved state, the rate of recovery from the starved state to the modified full state, and the maintenance rates of both, will be different, such that *σ′* = *σ*(*M*′), *ρ′* = *ρ*(*M*′), *β′* = *β*(*M*′), *δ′* = *δ*(*M*′). Rates of starvation and recovery for the invading population are easily derived by adjusting the starting or ending state before and after starvation and recovery, leading to the following timescales:

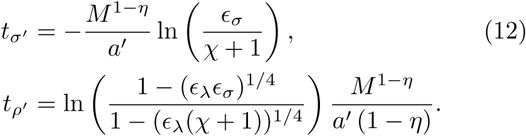

The maintenance rates for the invading population require more careful consideration. First, we must recalculate the yields *Y*, as they must now be integrated over life stages that have also been slightly modified by the addition or subtraction of body fat reserves. Given an energy density of *E_d_* = 18200 (J g^−1^) for grass [14] the maintenance value of the invading population is given by 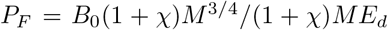, and the yield for a full organism will be given by *Y_F_* = (1 + *χ*)*M E_d_/B′_λ_* (g individual ⋅ g grass ^−1^) where 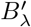 is the lifetime energy use for the invading population reaching maturity given by

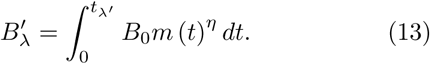

where

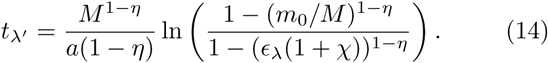

Note that we do not use this timescale to determine the reproductive rate of the invading consumer—which is assumed to remain the same as the resident population—but only to calulate the lifetime energy use. Similarly, the maintenance for hungry individuals 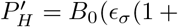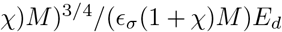 and the yield for hungry individuals (representing the cost on resources to return to the full state) is given by 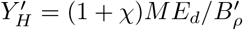 where

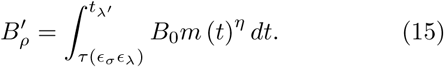

Finally, we can calculate the maintenance of the invaders

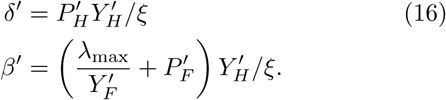

To determine whether or not the invader or resident population has an advantage, we compute *R^*^* (*M*) and *R^*^*(*M*′ = *M*(1 + *χ*)) for values of *χ* ∈ (−*f*_0_*M*^γ−1^, 1/∊λ − 1), and the invading population is assumed to have an advantage over the resident population if *R^*^*(*M*′) < *R^*^*(*M*).

## Sensitivity to additional death terms

**Figure S1:**
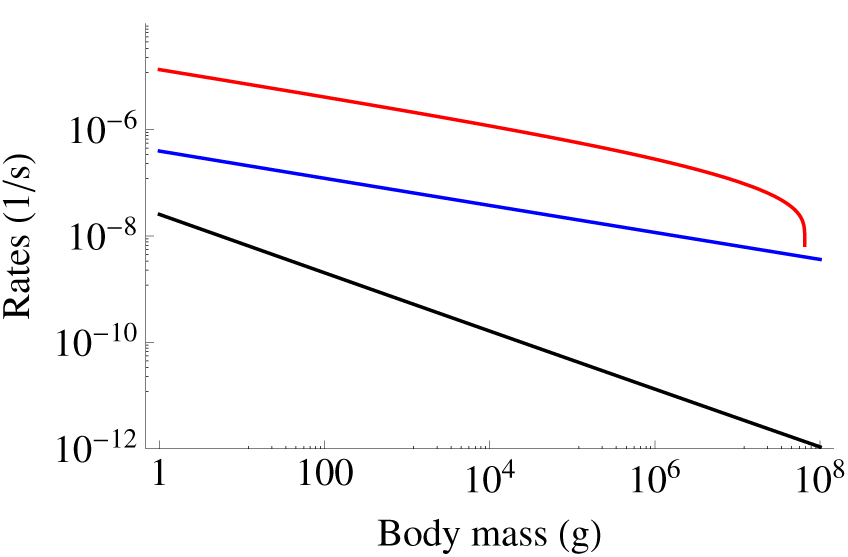
The rates of reproduction λ (blue), starvation-based mortality *μ* (red), and survivorship-based death 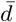 (black) as a function of adult mass.

It should be noted that our set of dynamics (Equations 7 and 9) could include a constant death term of the form *−d_F_F* and *−d_H_H* to represent death not directly linked to starvation. Adding terms of this form to our model would simply adjust the effective value of *λ* and *μ*, and we could rewrite Equation 9 with *λ′* = λ − *d* and *μ′* = *μ* − *d*. These substitutions would not alter the functional form of our model nor the steady-states and qualitative results, however the quantitative values could shift based on the size of *d* relative to *λ* and *μ*.

Survivorship has a well-known functional form which changes systematically with size (e.g. [17]). Typically survivorship is defined using the Gompertz curve

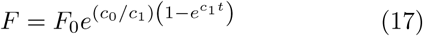

where the parameters have the following allometric dependencies on adult mass *c*_0_ = *a*_0_*M*^*b*_0_^ and *c*_1_ = *a*_1_*M*^*b*_l_^, with *a*_0_ = 1.88 × 10^−8^ (s g^−*b*_0_^), *b*_0_ = −0.56, *a*_1_ = 1.45 × 10^−7^ (s g^−*b*_l_^), and *b*_1_ = −0.27 (see [17] for a review).

We are interested in the specific death rate of the form 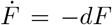 and using the derivative of Equation 17 we find that 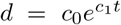. Our model considers the average rates over a population and lifecycle and the average death rate is given by

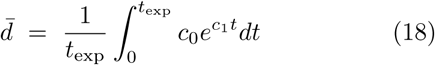

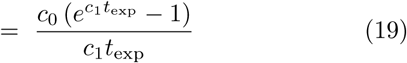

where *t*_exp_ is the expected lifespan following the allometry of *t*_exp_ = *a*_2_*M*^*b*_2_^ with *a*_2_ = 4.04 × 10^6^ (s g^−*b*_2_^) and *b*_2_ = 0.30 [17, 18]. Given the allometries above we have that

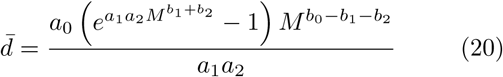

which scales roughly like *M*^*b*_0_^ because *b*_1_ and *b*_2_ are close in value but opposite in sign. In Figure SS1 we compare the value of 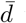 to the reproductive, λ, and starvation-based mortality, *μ*, rates. The values of 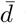 are orders of magnitude smaller than these rates for all mammalian masses, and thus, adding this non-starvation based death rate to our model does not shift our results within numerical confidence.

**Figure S2:**
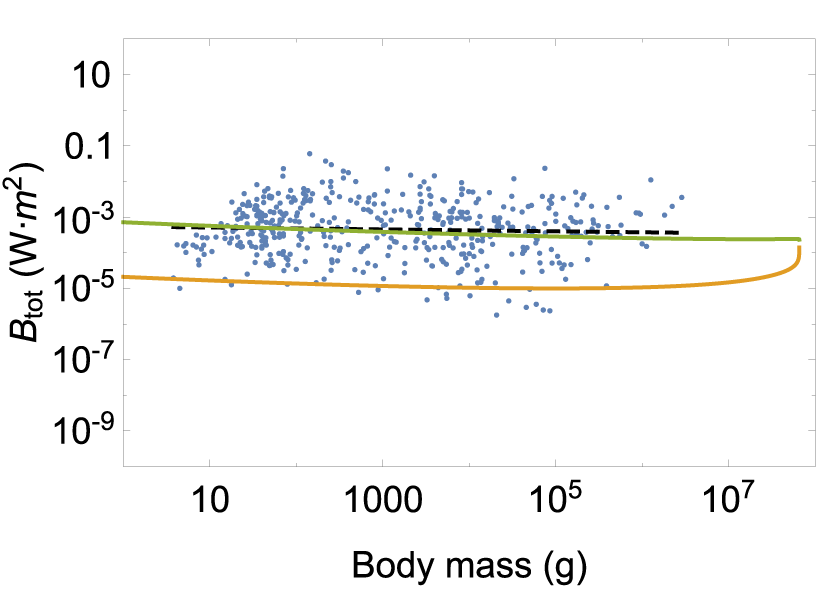
Total energetic use *B*_tot_ of consumer populations at the steady state as a function of body mass (*F^*^* is shown in green and *H^*^* in orange). The data are from Damuth [21] and have been converted to total population metabolism using the allometric relationships for metabolic rate (e.g. Refs. [3–5]).

## NSM and the energy equivalence hypothesis

The energy equivalence hypothesis is based on the observation that if one assumes that the total metabolism of an ecosystem *B*_tot_ is equally partitioned between all species (*B_i_*, the total metabolism of one species, is a constant), then the abundances should follow *N* (*M*) *B* (*M*) = *B_i_*, implying that *N* (*M*) ∝ *M^−η^*, where *η* is the metabolic scaling exponent [19, 20]. As *η* ≈ 3/4 this hypothesis is consistent with Damuth’s law [19]. However, the actual equivalence of energy usage of diverse species has not been measured at the population level for a variety of whole populations. Figure SS2 recasts the results of the NSM in terms of this hypothesis and shows that *F^*^B* is nearly constant over the same range of mammalian sizes up to the asymptotic behavior for the largest terrestrial mammals.

## Application of NSM limits to aquatic mammals

A theoretical upper bound on mammalian body size is given by *∊_σ_* = 0, where mammals are entirely composed of metabolic reserves, and this occurs at *M* = 8.3 × 10^8^ (g), or 120 times the mass of a male African elephant. We note this particular limit as it may have future relevance to considerations of the ultimate constraints on aquatic mammals.

